# Single-cell classification using learned cell phenotypes

**DOI:** 10.1101/2020.07.22.216002

**Authors:** Yang Chen, Tadepally Lakshmikanth, Jaromir Mikes, Petter Brodin

## Abstract

Single-cell methods such as flow cytometry, Mass cytometry and single-cell mRNA sequencing collect high-dimensional data on thousands to millions of individual cells. An important aim during the analysis of such data is to classify cells into known categories and cell types. One commonly used approach towards this is clustering of cells with similar features followed by manual annotation of clusters in relation to known biology. A second approach, commonly used for cytometry data relies on manual sorting or “gating” of cells, often based on pairwise combinations of measurements used in a stepwise and very tedious process of cell annotation. Both of these approaches require manual inspection and annotation of every new dataset generated, a process that is not only time consuming but also subjective and surely influential for the conclusions drawn. The manual annotation is also difficult to reproduce by other researchers with a different perception of features that signify their cells of interest. Here we propose an alternative strategy based on machine learning of known phenotypes from manually curated, high-dimensional data and thereby enabling rapid classification of subsequent datasets in a more reproducible manner. This simple approach increases both throughput, reproducibility and simplicity of cell classification in single-cell biology.

---

The number of methods for generating high-dimensional single-cell data is rapidly expanding. These methods are enabling multi-scale analyses from thousands to millions of cells, resulting in unprecedented levels of details in our analyses of heterogeneous tissues (Davis and Brodin, 2018a). (Davis and Brodin, 2018b). The most widely used single-cell technology to date has been flow cytometry, developed more than 50 years ago, and still a workhorse in the fields of immunology, cancer biology and cell biology. Flow cytometry allows for the rapid enumeration of close to 30 parameters in millions of cells using antibodies coupled to fluorescent probes. With the application of mass cytometry, the number of measured parameters in any given cell has increased to > 50 (Brodin, 2019). Single-cell genomic methods is also advancing rapidly and now allow for the quantification of thousands of mRNA-molecules, epigenetic states, or DNA-sequence variants, all in tens of thousands of cells (Macaulay et al., 2017). After normalization and various other steps of pre-processing, one important task in the analysis of such high-dimensional single-cell data, is to classify cells into defined populations that can be interrogated and interpreted in relation to what is already known with respect to known biological functions. In flow and mass cytometry, this process is most commonly done by manually inspecting individual markers and pairwise combinations of markers with known distribution, through an iterative process of “gating”, cells are manually classified into the many known populations of cells. For blood immune cell about 20 different cell lineages and many subpopulations have been described (Uhlen et al., 2019). This process is intractable when the number of measured features grows above a certain level, such as in mass cytometry and single-cell mRNA-sequencing datasets (Mair et al., 2016). This manual gating procedure must also be repeated in full, with every new sample, and every experiment. Given these well-known limitations, several methods for automated cell classification have been introduced in recent years (Mair et al., 2016). There is now a plethora of tools for clustering of single-cell data, and the specific tool used in the field largely depends on the type of data, and the habit of the investigator although some systematic comparisons have been performed (Weber and Robinson, 2016). One common feature of existing tools today, is the requirement for clusters of cells, like manually gated cell populations, to be curated by the investigator in analyzing each new dataset. This represents another laborious and subjective process, that strongly influences the downstream conclusions. The subjective nature of this cluster annotation also makes it difficult to document, and therefor also difficult to reproduce, by other investigators. Even if complete misclassification of cells is probably uncommon in these instances, subtle variations in the interpretation of cell phenotypes will occur that make comparisons across studies difficult. To tackle these challenges, we propose an alternative strategy, in which cell classification is manually performed, based on expert knowledge, only during the assembly of a reference dataset, and this dataset is then used to train a classifier algorithm that will be able to classify similar cells rapidly and with good accuracy in additional datasets generated under comparable conditions. This approach increases the speed at which cell classification can occur, but also makes the process more reproducible.

We use large scale mass cytometry datasets to illustrate our approach, although we anticipate this strategy will be equally useful for other single-cell datasets. We first built a software for gating training datasets, not based on traditional biaxial plots as is commonly done during flow cytometry “gating”. Instead we embedded a dataset using Barnes-Hut-SNE (Maaten, 2013) and gated training data based on all relevant antigens in the panel. In the first step, we embed an entire whole blood dataset of ~1,000,000 cells using bhSNE and color the data by canonical lineage-defining markers such as CD3, CD4, CD8, CD14, CD56 and CD19, until all subpopulations have been identified and manually gated in the bh-SNE embedded data. Larger clusters, such as CD4^+^ T-cells, CD8^+^ T-cells, B-cells, NK-cells, and Monocytes are subsequently embedded separately in a second level bh-SNE embedding, taking only features relevant for the given cell type into account. The purpose of this second layer of selection is to identify and label subpopulations of cells of importance (Maecker et al., 2012), and include these when training the learning algorithm (Figure 1). Examples of such subpopulations identified in this second level classification are memory and naïve lymphocyte subsets, and canonical subpopulations of monocytes and NK-cells (Figure 1A). It is important to note that these subpopulations can be more difficult to discriminate at the first level bhSNE embedding where all cell lineages are included. Other methods for generating labeled training data, such as manual flow cytometry gating and supervised or unsupervised clustering of cells can also be used for the following training algorithm.

**Figure 1.**
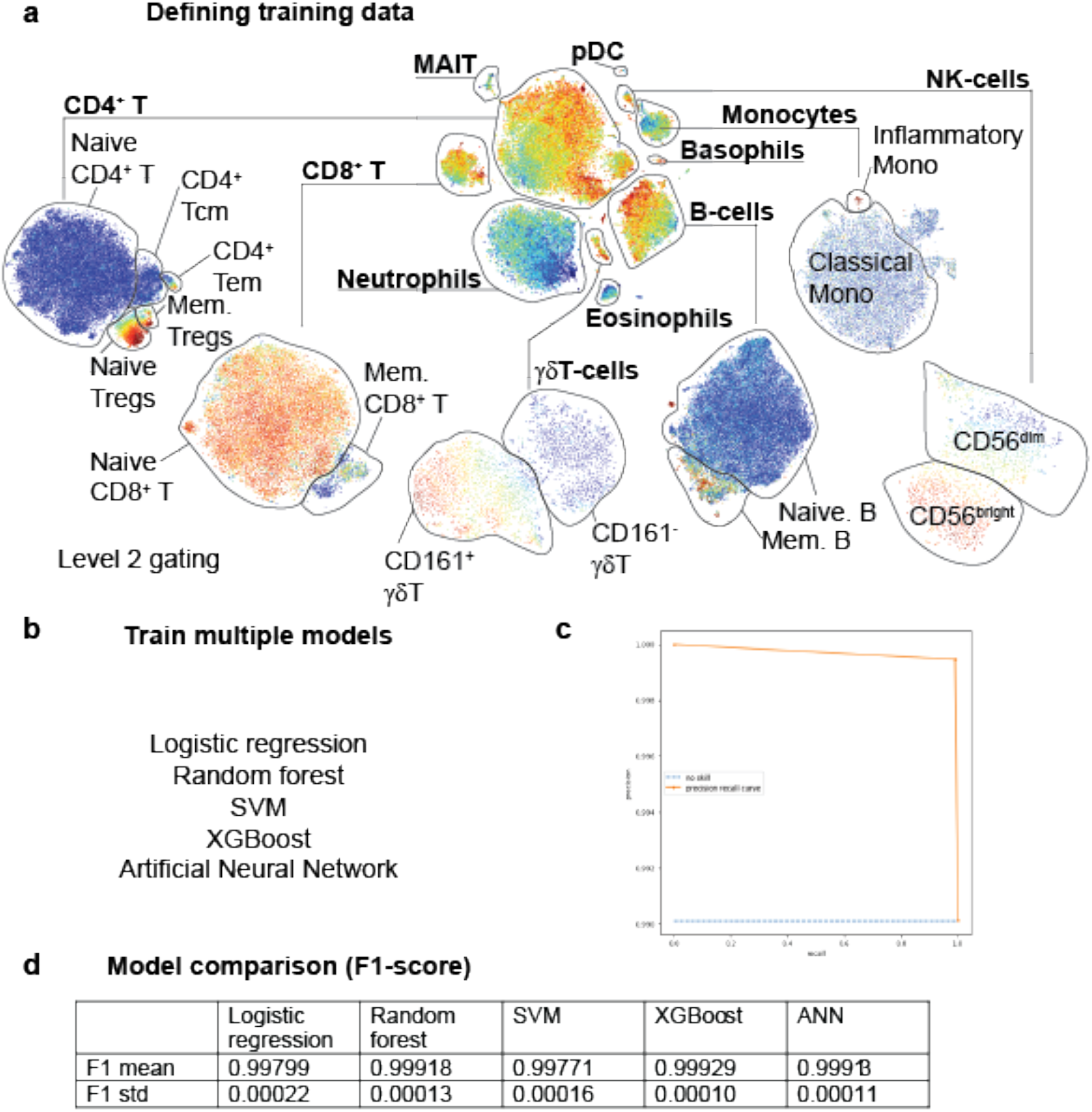
**a**) Manual gating using a hierarchical tSNE embedded mass cytometry data. The indicated example markers are highlighted in color from blue (low) to red (high). **b**) The classified training data is used to train different learning algorithms. **c**) Precision/recall curves. **d**) F1-score comparison of trained models.

After the training dataset of high quality has been classified and single-cell tables have been appended with cell labels, we are ready to train a classifier algorithm. The training process employs 4 common machine learning algorithms; Logistic regression (Yu et al., 2011), Random forest (Ho, 1998), support-vector machine (SVM)(Cortes and Vapnik, 1995), Extreme gradient boosting (XGBoost)(Friedman, 2001) Figure 1b). For each of these algorithms, the F1-score is calculated from precision and recall tests against an additional manually classified dataset not included during the training procedure (Figure 1c). From the mean and standard deviation of the F1-score, the best performing model is selected and used henceforth.

**Particles - Level 0**

| Population | F1 score |
| --- | --- |
| Cells | 0.99901 |
| Non-cells | 0.93633 |

**Whole blood - Level 1**

| Population | F1 score |
| --- | --- |
| B | 0.99576 |
| Basophils | 0.99509 |
| CD4T | 0.99458 |
| CD8T | 0.98649 |
| Eosinophils | 0.99377 |
| gdT | 0.97842 |
| MAIT | 0.92922 |
| Monocytes | 0.98639 |
| Neutrophils | 0.99740 |
| NK | 0.98161 |
| pDC | 0.98293 |

**CD4^+^ T-cells - Level 2**

| Population | F1 score |
| --- | --- |
| Central memory CD4T | 0.82282 |
| Effector memory CD4T | 0.85544 |
| Memory Tregs | 0.83341 |
| Naïve CD4T | 0.98413 |
| Naïve Tregs | 0.92499 |
| TEMRA CD4T | 0.55489 |

B-cells – Level 2

| Population | F1 score |
| --- | --- |
| IgD<sup>+</sup> Memory B | 0.74591 |
| IgD<sup>-</sup> memory B | 0.88233 |
| Naive B | 0.98359 |
| Plasmablasts | 0.06452 |
| Transitional B | 0.48740 |

NK-cells – Level 2

| Population | F1 score |
| --- | --- |
| CD56bright NK | 0.95993 |
| CD56dim NK | 0.97871 |

CD8^+^ T-cells – Level 2

| Population | F1 score |
| --- | --- |
| Activated CD8 T | 0.70737 |
| Central memory CD8 T | 0.73160 |
| DP T-cells | 0.90411 |
| Effector memory CD8T | 0.81967 |
| Naive CD8 T | 0.98115 |

Monocytes – Level 2

| Population | F1 score |
| --- | --- |
| Classical Mono | 0.98874 |
| Non-classical Mono | 0.83288 |
| Proinflammatory mono | 0.93532 |

γδT-cells – Level 2

$\gamma\delta$ T-cells – Level 2
| Population | F1 score |
| --- | --- |
| CD161<sup>+</sup> gdT | 0.98340 |
| CD161<sup>-</sup> gdT | 0.95848 |

The classification of subsequently generated datasets can now be performed in a fully automated fashion. Classifier finishes 1,000,000 cells in 36 seconds, and we recently classified a large experiment with 400 large FCS-files in 48 minutes while manual gating of comparable 34 subsets would take weeks to finish. It is important to note that the performance of the classifier must be investigated by visualization of the data output to ensure correct cell labeling. This is particularly important when test data is different from the training data as this obviously increases the uncertainty in the classification. We therefore redo training with the inclusion of new training data on a regular basis and redo the learning process.

Another important aspect of this approach is the ability to assess feature importance for the classification accuracy. This analysis can help guide better panel design in flow and mass cytometry in a data-driven manner to maximize the information content in a panel. This is done by iteratively applying the XGBoost classifier on the manually classified test dataset. In each loop one marker is excluded and the accuracy of the classification is monitored. From loss of precision, a score of importance can be assigned to each feature in the dataset (**Figure 2a**). The results of this exercise are shown for level 1 classification of 11 canonical blood immune cell populations and reveals that the most important marker is CD38, followed by CD99. We noticed that lineage markers such as CD19, CD20, CD20 and CD8a were absent from the list of the top 10 most important markers. This illustrates the ability of a data-driven approach such as this machine learning approach to guide better panel design in mass cytometry and flow cytometry. Similarly, we assess marker importance at each of the level 2 learning steps and report these in **Figure 2c**. We conclude from these findings that a machine learning approach weighting 4 common classification algorithms, represents an efficient method for automating cell classification by considering manually classified training data. The approach is fully reproducible, and the steps of cell classification are traceable when applied to novel data. The method also represents a massive increase in throughput for mass cytometry data analysis over manual gating, with hundreds of files readily classified in only a few seconds.

**Figure 2.**
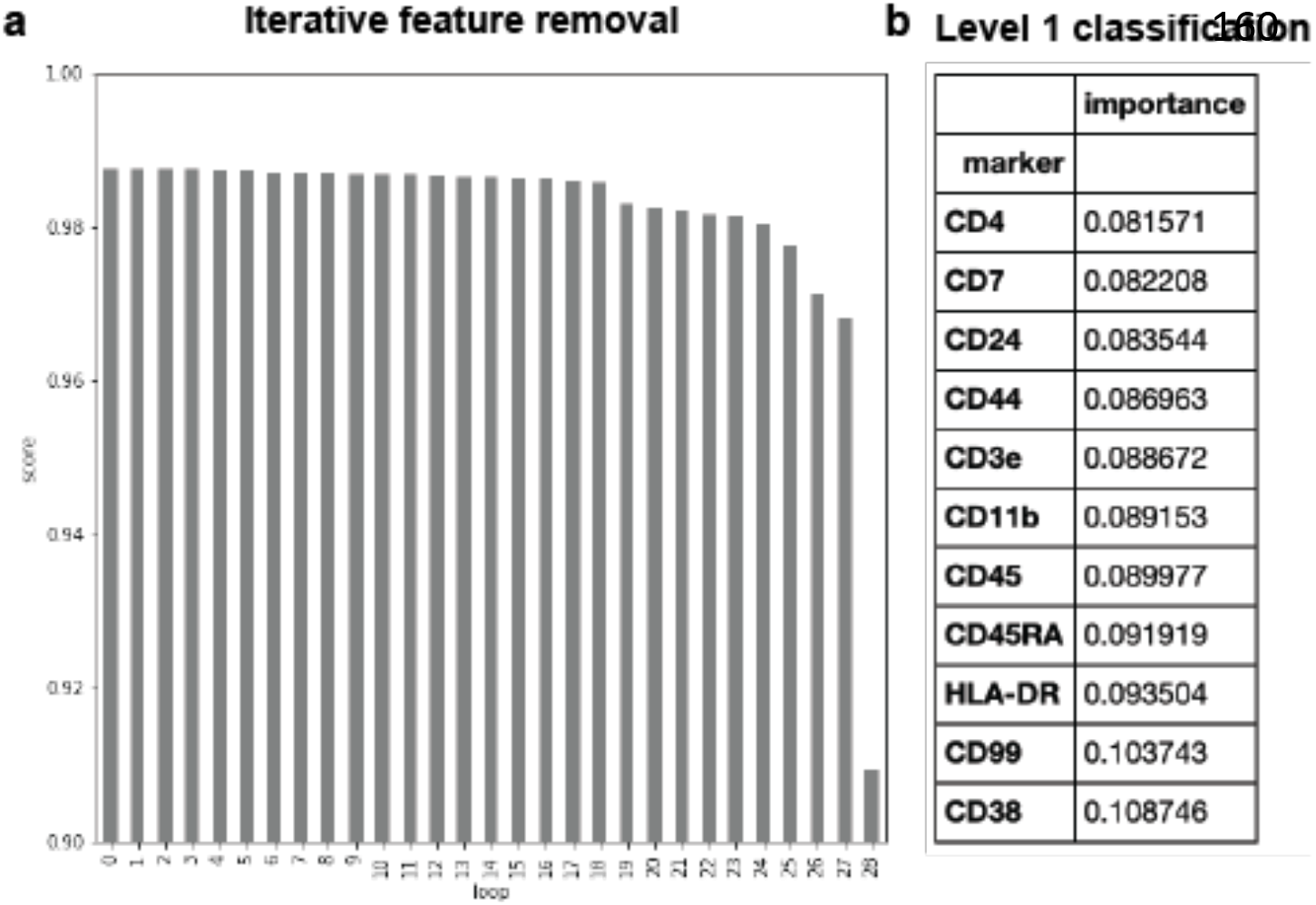
**a**) Iterative leave-one-marker out test of model performance reveal relative importance of each marker in the mass cytometry antibody panel. **b**) at level 1 cell populations classification top 11 markers are indicated for the example dataset classified in figure 1.

There is a need to improve both the throughput and reproducibility of classification of cell populations from single-cell data. Such datasets are now very commonly used but the conclusions drawn about biological processes, differences in cell composition and function in relation to health and disease, all depend upon the annotation of individual cells into canonical or novel cell populations. Manual processes have dominated in the past, particularly when flow cytometry datasets are used. Manual gating has several important caveats. It is intractable for datasets with hundreds of samples, millions of cells and tens to hundreds of features measured per cell. Also, manual annotation is subjective and difficult to reproduce and thereby influence the conclusions drawn from the data. The benefit of the manual and supervised analyses methods are that prior knowledge allow for rapid quality control of the data, there is a large amount of human experience involved in looking at the data which allow for biological interpretation and hypothesis formulation, not possible by any algorithms.

Here we present a very simple, yet powerful method for combining the interpretability of manual annotation, with the scalability, throughput, speed and reproducibility that automated methods allow. By using a manually annotated training dataset and learning cell phenotypes associated with annotations such as immune cell subsets, we are able to rapidly classify and annotate new datasets generated and still be able to draw biological conclusions from the data and relate these immediately to existing body of literature. A second important feature of this approach is that markers used for classification can be ranked by importance and thereby allow for optimization of antibody panels or mRNA-probes or other predefined panels of features analyzed. We provide the learning and classifier algorithms as a standalone software package for others to use freely. This software takes a defined input format and learns indicated cell population labels for future classification tasks. The main limitation of this work is that it is intrinsically difficult to develop a learning algorithm that performs equally well on all populations of interest. We typically find that small subsets require more training data and that subsets of cells without clear separation are obviously harder. In some instances, we find it useful to classify the well separated subsets and then within such subsets where phenotypes vary along a continuum, we apply one of several trajectory inference methods or dimensionality reduction approaches rather than assessing discrete subpopulations.

## Software availability

The method is implemented in Python and can be installed via pip. For the documentation, refer to the package repository at https://github.com/Brodinlab/cellgrid.

## Acknowledgements

P.B and Y.C designed and conceptualized the method. Y.C wrote all the code and implemented the methods. T.L generated the mass cytometry data.

